# XhetRel: A Pipeline for X Heterozygosity and Relatedness Analysis in Sequencing Data

**DOI:** 10.1101/2024.12.26.627867

**Authors:** Barış Salman, Nerses Bebek, Sibel Uğur İşeri

## Abstract

**Motivation:** Checking sample sex is one of the preliminary controls performed in genetic studies. Comparing multiple individuals and families in terms of sex and relatedness allows us to eliminate systematic errors in downstream analyses. While developing the pipeline we further delved into the origins of X heterozygosity by analysing the chromosome X variants from the 1000 Genomes Project.

**Result:** We created an accessible and user-friendly notebook application for X heterozygosity analysis. XhetRel can serve as an initial quality control step in sequencing projects.

Further investigating the source of X heterozygosity reveals the limitations of sequencing and variant calling methods in heterozygosity analysis. Our findings point to specific pseudogenes and gene clusters, such as *SLC25A5* and *GAGE* cluster, as key contributors to erroneous variant allele fractions.

**Availability and implementation:** Source code is available https://github.com/barslmn/XhetRel.

Collab notebook can be accessed at https://colab.research.google.com/drive/1ep69JvXLwK5ndHUQ8qIGTWvauzsTW9fi.

## Introduction

X and Y chromosomes originate from a shared ancestral autosomal pair. (Lahn & Page, 1999) Over the course of molecular evolution, these chromosomes have accumulated repetitive and highly homologous regions, making them challenging to sequence analyses. (Pinto et al., 2023) Despite efforts to address this issue, current bioinformatics methods often exclude the sex chromosomes due to their biological variability. (Webster et al., 2019)

The allele frequencies and heterozygous variants on the X chromosome in male individuals are often erroneous. (Wang et al., 2022) While new strategies have been developed to address these biological challenges, widespread approaches that rely on a diploid model for variant calling in sex chromosomes continue to produce these errors.

During exome sequencing analyses, heterozygous variants on the X chromosome are often observed in male samples. However, this does not accurately reflect biological status, as males carry only one X and are hemizygous for both sex chromosomes. The exception lies in the pseudoautosomal regions (PARs), located at the terminal regions of the X and Y chromosomes, which span approximately 3Mb together. These regions exhibit homology between the chromosomes, allowing males to be “homozygous” or “heterozygous” in these regions. This homology enables chromosomes to pair during cell division and facilitates recombination, which is restricted to PARs. (Raudsepp et al., 2012)

The heterozygous variants observed along the X chromosome excluding the PARs present discrepancy in males. In exome sequencing, such heterozygous variants in males can serve as a useful quality metric to predict the individual's sex.

Wrong sample records can arise from several factors, including incorrect self-declaration; errors in record-keeping or labeling, sample mix-ups, cross-contamination during downstream analyses, or sex chromosome aneuploidies.

We first applied this method after encountering the study by Do et al., who utilized it to verify sample sex as a part of quality control. (Do et al., 2015; Akçakaya et al., 2019) Combined with relatedness analysis, this approach can be a powerful tool for identifying sample mix-ups especially in studies involving multiple families.

Herein, we developed a user-friendly notebook and workflow designed to detect relatedness and predict sex based on X heterozygosity from large sequencing variant call format (VCF) files. Additionally, we empirically investigated the sources of X heterozygosity using data from the 1000 Genomes Project (1KGP). (Auton et al., 2015)

## Material and Methods

For relatedness analysis, VCF files are first filtered individually for allele fraction greater than 0.25 and read depth higher than 20, and merged into a single file, any variants with missing genotype fraction less than 20% further filtered out. (Figure 1A) The robust relationship inference method implemented in VCFtools is used for the relatedness. (Danecek et al., 2011) X chromosome non-PAR variants are queried using BCFtools, counting only variants with genotypes 0/1 and 1/1 (Danecek et al., 2021). X heterozygosity is calculated as the ratio of heterozygous variants to the total number of variants on chromosome X using AWK. The results are visualized with a custom plot generated in MultiQC. (Ewels et al., 2016) The pipeline was tested using two separate sons, father, mother trios of Ashkenazi Jewish and Han Chinese ancestry from The Genome in a Bottle Consortium.(*Genome-in-a-Bottle/Giab_data_indexes: This Repository Contains Data Indexes from NIST's Genome in a Bottle Project*., n.d.; Zook et al., 2016)(https://github.com/genome-in-a-bottle/giab_latest_release) (“Genome in a Bottle—a Human DNA Standard,” 2015) We have combined these to approaches with a tool named Xhetrel.

**Figure 1:**
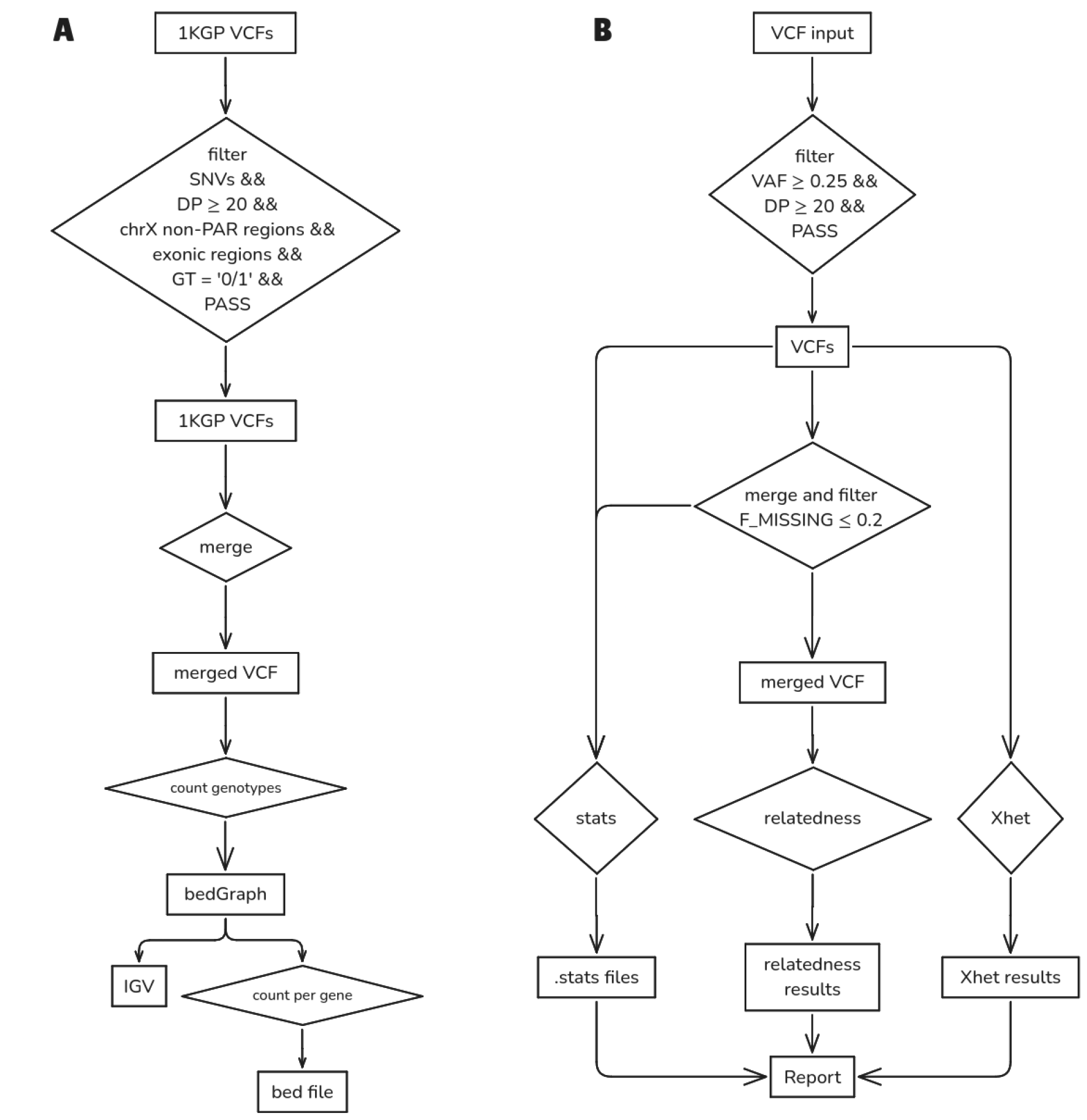
Workflow diagrams. A) Workflow diagram for XhetRel tool. B) Workflow diagram for 1000 genomes project analyses.

To investigate the sources of the heterozygosity, we analysed X chromosome genotypes from 1,223 male samples in the 1000 Genomes Project. Variant calls are filtered for single nucleotide variants (SNVs) in exonic regions using three variant allele fraction (VAF) cut-offs: (i)25 ≤ VAF ≤ 75, (ii)33 ≤ VAF ≤ 66, (iii)45 ≤ VAF ≤ 55. The number of male individuals with variants in each region was counted, and the top 20 regions were annotated. (Figure 1B)

Information on paralogous genes was obtained from the Ensembl Comparative Genomics tab, while the list of pseudogenes for a given gene was retrieved from the HUGO Gene Nomenclature Committee (HGNC) database (https://genenames.org). Pseudogenes are identified using UCSC BLAT (https://genome.ucsc.edu/cgi-bin/hgBlat).

## Results

### XhetRel tool

We have developed a preliminary quality control workflow process for sex and relatedness. The workflow is available as an online Colab notebook or can be executed locally using Nextflow.

The relatedness analysis results are visualized in Figure 2. The heatmap (Figure 2A) displays kinship coefficients (*ϕ*), where values of approximately 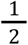 indicate monozygotic twins or self-comparisons, and values of approximately 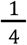 indicate first-degree relationships such as parent-offspring pairs. (Manichaikul et al., 2010) The sample are notated as follows: 'A' and 'C' denote Ashkenazi Jewish and Chinese trios respectively, while 'i', 'f', and 'm' represent index (child), father, and mother, respectively. Figure 2B illustrates X chromosome heterozygosity patterns, with maternal samples showing expected heterozygosity rates around 0.5, while paternal and male offspring samples demonstrate rates approaching zero, consistent with their hemizygous state for the X chromosome.

**Figure 2:**
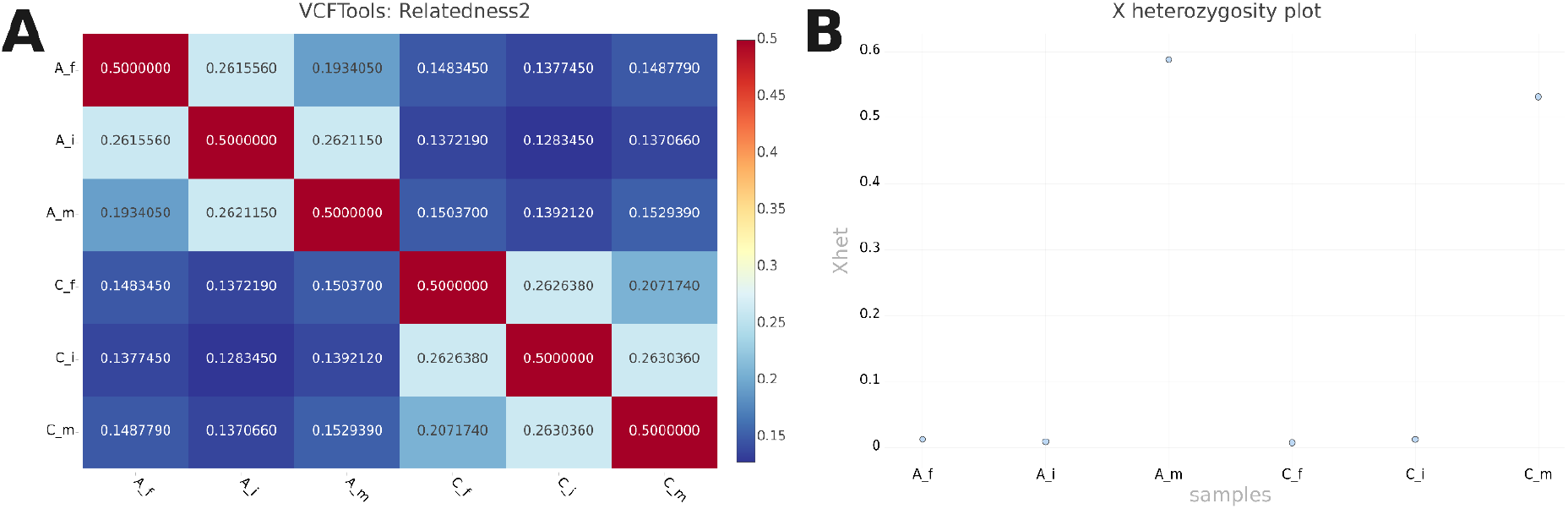
Relatedness and X heterozygosity plots generated by MultiQC. A) Heatmap displaying individual relatedness. A: Individual from the Ashkenazi Jewish trio; C: Individual from the Chinese trio; f: father; m: mother; i: index. B) Custom plot illustrating the fraction of heterozygous variants on the X chromosome.

### 1000 Genome Project analyses

We have plotted the heterozygous regions using IGV with the major evolutionary X regions and minor ampliconic and low complexity regions. For the VAF cut-off between 0.25 and 0.75, X-conserved regions carry the most of the heterozygous variation in major regions with density of variation per megabase being 194.4, 433.2, 2.3, 789 for X-added region (XAR), X-conserved region 1 (XCR), X-transposed region (XTR), and XCR2 respectively. Ampliconic regions 5 and 9 were the highest amount minor regions with the density 6945.5, and 1013.2 respectively. (Figure 3)

**Figure 3:**
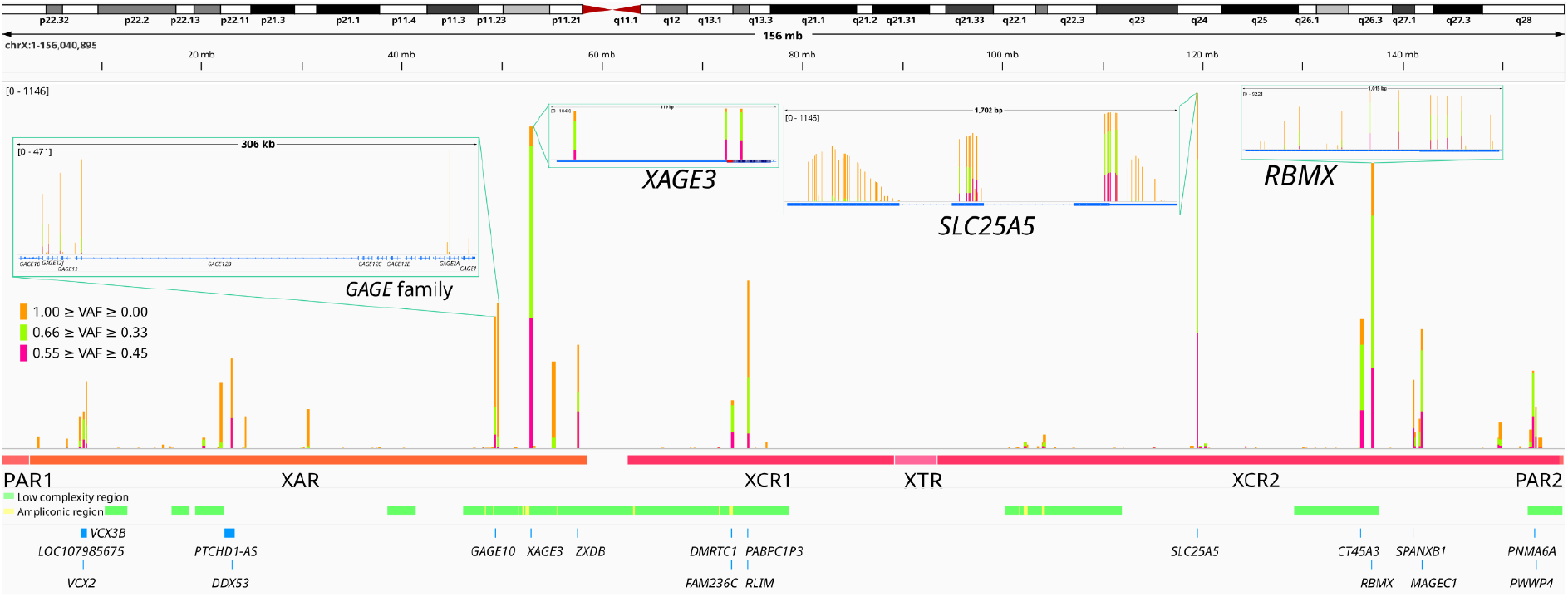
Genotype counts across chromosome X for alternative allele fraction cut-offs. Low complexity and ampliconic regions were identified based on the list from Cotter et al., 2016, while major chromosome X regions were defined using the 2019 Webster et al. study. (Cotter et al., 2016; Webster et al., 2019) PAR: pseudoautosomal region, XAR: X-added region, XCR: X-conserved region, XTR: X-transposed region.

A total of 71,553, 27,504 and 9,243 heterozygous genotype calls were identified across in 1,223 male samples, corresponding to the applied VAF cut-offs. Table 1 lists genotype counts for top 20 genes.

**Table 1.**
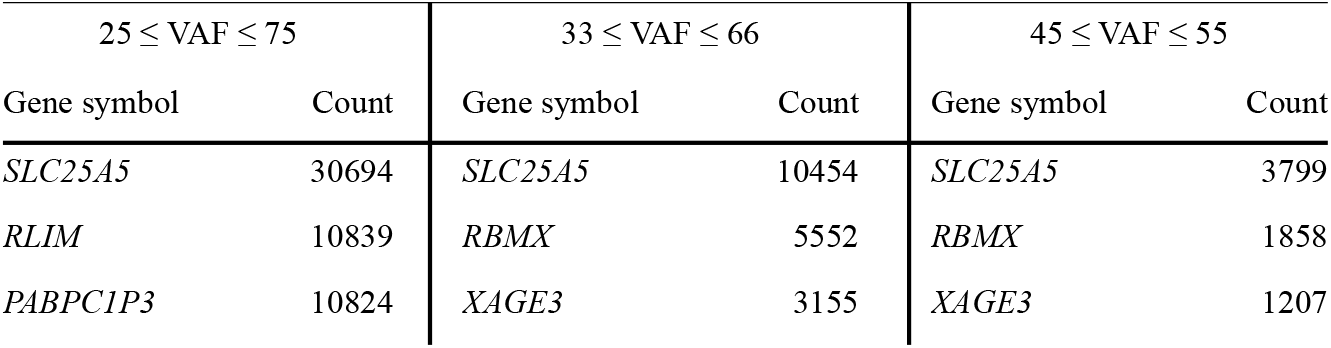
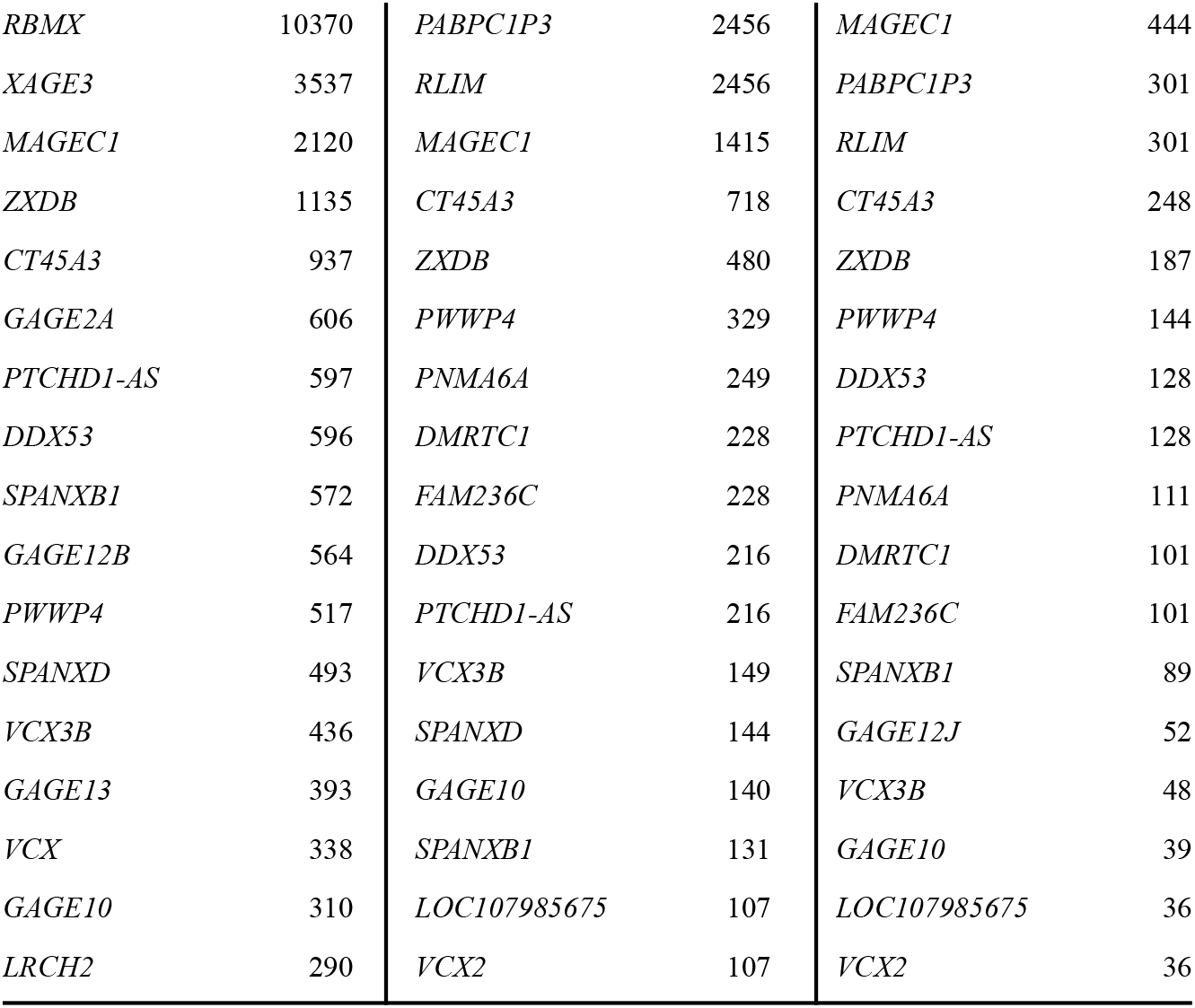
Genotype counts for top 20 genes for alternative VAF cut-offs.

The *SLC25A5* gene consistently ranked as the top gene across all VAF cut-offs, with nearly 3,800 heterozygous calls even under the most stringent VAF cut-off (45 ≤ VAF ≤ 55). To investigate this further, we conducted detailed analysis using UCSC-BLAT against hg38 reference genome on the *SLC25A5* mRNA sequence (1,307 bases: NM_001152.5/ENST00000317881.9). Results were filtered for matches with a percent identity greater than 80% and alignment span size exceeding 1,000 bases. The analysis revealed strong matches across 10 autosomal loci, including chromosomes 2, 4, 5, 6, 7, 9, 13, and 22. Additionally, a homologous region between the X and Y chromosomes showed a relatively lower degree of match, with 81.20% identity over a 4,712-base span (Table 2).

**Table 2.** UCSC-BLAT Analysis of the *SLC25A5* mRNA Sequence. Matches are based on UCSC-BLAT scores against the hg38 reference genome: first row is the query sequence match, there are nine hits on autosomal loci involving pseudogenes, one hit on SLC24A4, and two hits on gonadal chromosomes (excluding the query locus).

| Query | Start | End | Identity | Strand | Chr | Start (bp) | End (bp) | Span Size (bp) | Gene Included |
| --- | --- | --- | --- | --- | --- | --- | --- | --- | --- |
| NM_001152.5 | 1 | 1307 | 100.00% | F | chrX | 119,468,444 | 119,471,396 | 2,953 | *SLC25A5* |
| NM_001152.5 | 1 | 1227 | 93.90% | F | chr7 | 32,471,429 | 32,472,998 | 1,570 | *SLC25A5P5* |
| NM_001152.5 | 5 | 1234 | 93.90% | F | chr7 | 54,419,371 | 54,420,600 | 1,230 | *SLC25A5P3* |
| NM_001152.5 | 6 | 1222 | 92.90% | R | chr22 | 42,000,445 | 42,002,033 | 1,589 | *SLC25A5P1* |
| NM_001152.5 | 14 | 1223 | 92.90% | R | chr2 | 33,839,529 | 33,840,742 | 1,214 | *SLC25A5P2* |
| NM_001152.5 | 2 | 1203 | 92.30% | F | chr6 | 121,653,739 | 121,654,924 | 1,186 | *SLC25A5P7* |
| NM_001152.5 | 1 | 1230 | 92.00% | R | chr9 | 32,333,195 | 32,334,431 | 1,237 | *SLC25A5P8* |
| NM_001152.5 | 14 | 1230 | 91.30% | R | chr4 | 186,328,455 | 186,329,663 | 1,209 | *SLC25A5P6* |
| NM_001152.5 | 48 | 1179 | 88.80% | F | chr13 | 57,314,711 | 57,316,417 | 1,707 | *SLC25A5P4* |
| NM_001152.5 | 100 | 347 | 87.70% | F | chr4 | 185,143,400 | 185,144,927 | 1,528 | *SLC25A4* |
| NM_001152.5 | 58 | 1224 | 85.10% | R | chr5 | 75,752,294 | 75,753,498 | 1,205 | *SLC25A5P9* |
| NM_001152.5 | 91 | 810 | 81.20% | R | chrY | 1,387,280 | 1,391,991 | 4,712 | *SLC25A6* |
| NM_001152.5 | 91 | 810 | 81.20% | R | chrX | 1,387,280 | 1,391,991 | 4,712 | *SLC25A6* |

These 12 loci was then annotated for their involved genes. Nine of 10 autosomal loci corresponded to an individual *SLC25A5* pseudogenes (*SLC25A5P1-9*). These pseudogenes are classified as transcribed processed or processed-only pseudogenes, indicating multiple retrotransposition events during genome evolution. Furthermore, S*LC24A4* on chromosome 4 and *SLC25A6* on chromosomes X and Y are paralogue genes for *SLC25A5*. These findings highlight extensive homology of SLC24A5 across multiple chromosomes, posing significant challenges for short-read next generation sequencing approaches and resulting in ambiguous outcomes.

The second factor contributing to male X chromosome heterozygosity involves the *RBMX* gene, which has 36 paralogues and 5 pseudogenes (*RBMXP1*–*5*). Among these, the paralogue *RBMXL1*, located on chromosome 1, exhibits a striking similarity to the X-linked *RBMX* gene, with 99% identity in cDNA alignment and 95% identity across the entire gene.

Although the *RBMX* gene lacks a direct gametolog on the Y chromosome, several Y-linked paralogues—*RBMY1A, RBMY1B, RBMY1D, RBMY1E, RBMY1F*, and *RBMY1J*—have been identified, each sharing approximately 50% identity across their full gene sequences.

As mentioned earlier for *SLC25A5*, this extensive network of paralogous genes, combined with the presence of five pseudogenes, likely accounts for the inflated male heterozygosity observed for *RBMX* in whole-exome sequencing (WES) studies.

Another notably contributor to the X heterozygosity is the GAGE family. GAGE family is 16 gene cluster residing in p11.23 on chromosome X with high sequence identity. The gene cluster is a result of a recent evolutionary event which found in primates.(Liu et al., 2008) All the genes have high sequence identity and contains L1 except for *GAGE1* which is considered to be the ancestral gene. (Gjerstorff & Ditzel, 2008)

## Discussion

### XhetRel pipeline

We have developed a versatile pipeline that can be run directly on a Google Colab notebook without requiring any local installations. Alternatively, it can be integrated into larger sequence analysis workflows. The reported metrics enable the identification of major sample mix-ups prior to downstream variant analysis, ensuring data integrity.

While the pipeline is user-friendly and accessible on Google Colab, larger datasets may benefit from being processed locally using Nextflow and Docker for improved performance and scalability.

It is important to note that relatedness information derived from the relatedness2 package cannot be utilized as evidence for ACMG De Novo classifications. This limitation arises because the does not perform Mendelian check and in terms of relatedness, mother and female children; father and male children cannot be distinguished from each other.

### 1000 genomes project analyses

We have investigated male samples from 1000 genomes project to identify and analyse the origins of apparent heterozygous variants on the X chromosome, revealing that these variants primarily arise from pseudogenes of *SLC25A5*, paralogues genes of *RBMX* and highly homologous gene cluster the *GAGE* family.

Our genome has repetitive sequences scattered due to events like retrotransposition, duplications, and translocations. We can't separate where these reads are coming from and seeing them mistakenly aligned. In this study, we focused exclusively on chromosome X, considering the repetitiveness of the human genome the same errors can be expected all around the genome.(Hoyt et al., 2022) There will be other regions with false mappings and sequence errors in the genome that we will inadvertently see as heterozygous. We are just not looking because they're diploid regions and seeing heterozygous variants is expected. Because males only carry a single copy of chromosome X, that makes it a suitable model.

This discrepancy in variant fractions investigated with T2T genome as reference and long read sequencing which is a more promising technology for repetitive regions. (Miga et al., 2020) The X chromosome can be used to model *pseudogene and gene family — variant allele fraction* relation to recalibrate variant heterozygosity and to eliminate pseudo-heterozygosity of the variant calls in sequencing studies.

## Acknowledgements

Infrastructure supports for this study were provided by the NoroDIAB project (Istanbul University Scientific Research Fund No. 2019K12-149071) and the ISTisNA project (ISTKA No. TR10/22/THN). We also thank the Turkish Academy of Sciences for the 2019 Distinguished Young Scientist Award to SAUI.

## References

Akçakaya, N. H., Salman, B., Görmez, Z., Tarkan Argüden, Y., Çırakoğlu, A., Çakmur, R., Dönmez çolakoğlu, B., Hacıhanefioğlu, S., Özbek, U., Yapıcı, Z., & Uğur İşeri, S. A. (2019). A Novel and Mosaic WDR45 Nonsense Variant Causes Beta-Propeller Protein-Associated Neurodegeneration Identified Through Whole Exome Sequencing and X chromosome Heterozygosity Analysis. Neuromolecular Medicine, 21(1), 54–59. 10.1007/s12017-018-08522-6

Auton, A., Abecasis, G. R., Altshuler, D. M., Durbin, R. M., Abecasis, G. R., Bentley, D. R., Chakravarti, A., Clark, A. G., Donnelly, P., Eichler, E. E., Flicek, P., Gabriel, S. B., Gibbs, R. A., Green, E. D., Hurles, M. E., Knoppers, B. M., Korbel, J. O., Lander, E. S., Lee, C., … National Eye Institute, N. (2015). A global reference for human genetic variation. Nature, 526(7571), 68–74. 10.1038/nature15393

Cotter, D. J., Brotman, S. M., & Sayres, M. A. W. (2016). Genetic Diversity on the Human X Chromosome Does Not Support a Strict Pseudoautosomal Boundary. Genetics, 203(1), 485. 10.1534/genetics.114.172692

Danecek, P., Auton, A., Abecasis, G., Albers, C. A., Banks, E., DePristo, M. A., Handsaker, R. E., Lunter, G., Marth, G. T., Sherry, S. T., McVean, G., Durbin, R., & 1000 Genomes Project Analysis Group. (2011). The variant call format and VCFtools. Bioinformatics, 27(15), 2156–2158. 10.1093/bioinformatics/btr330

Danecek, P., Bonfield, J. K., Liddle, J., Marshall, J., Ohan, V., Pollard, M. O., Whitwham, A., Keane, T., McCarthy, S. A., Davies, R. M., & Li, H. (2021). Twelve years of SAMtools and BCFtools. GigaScience, 10(2), giab008. 10.1093/gigascience/giab008

Do, R., Stitziel, N. O., Won, H.-H., Jørgensen, A. B., Duga, S., Angelica Merlini, P., Kiezun, A., Farrall, M., Goel, A., Zuk, O., Guella, I., Asselta, R., Lange, L. A., Peloso, G. M., Auer, P. L., NHLBI Exome Sequencing Project, Girelli, D., Martinelli, N., Farlow, D. N., … Kathiresan, S. (2015). Exome sequencing identifies rare LDLR and APOA5 alleles conferring risk for myocardial infarction. Nature, 518(7537), 102–106. 10.1038/nature13917

Ewels, P., Magnusson, M., Lundin, S., & Käller, M. (2016). MultiQC: Summarize analysis results for multiple tools and samples in a single report. Bioinformatics (Oxford, England), 32(19), 3047–3048. 10.1093/bioinformatics/btw354

Genome in a bottle—A human DNA standard. (2015). Nature Biotechnology, 33(7), 675–675. 10.1038/nbt0715-675a

genome-in-a-bottle/giab_data_indexes: This repository contains data indexes from NIST's Genome in a Bottle project. (n.d.). Retrieved December 15, 2024, from https://github.com/genome-in-a-bottle/giab_data_indexes

Gjerstorff, M. F., & Ditzel, H. J. (2008). An overview of the GAGE cancer/testis antigen family with the inclusion of newly identified members. Tissue Antigens, 71(3), 187–192. 10.1111/j.1399-0039.2007.00997.x

Hoyt, S. J., Storer, J. M., Hartley, G. A., Grady, P. G. S., Gershman, A., de Lima, L. G., Limouse, C., Halabian, R., Wojenski, L., Rodriguez, M., Altemose, N., Rhie, A., Core, L. J., Gerton, J. L., Makalowski, W., Olson, D., Rosen, J., Smit, A. F. A., Straight, A. F., … O'Neill, R. J. (2022). From telomere to telomere: The transcriptional and epigenetic state of human repeat elements. Science, 376(6588), eabk3112. 10.1126/science.abk3112

Lahn, B. T., & Page, D. C. (1999). Four Evolutionary Strata on the Human X Chromosome. Science, 286(5441), 964–967. 10.1126/science.286.5441.964

Liu, Y., Zhu, Q., & Zhu, N. (2008). Recent duplication and positive selection of the GAGE gene family. Genetica, 133(1), 31–35. 10.1007/s10709-007-9179-9

Manichaikul, A., Mychaleckyj, J. C., Rich, S. S., Daly, K., Sale, M., & Chen, W.-M. (2010). Robust relationship inference in genome-wide association studies. Bioinformatics, 26(22), 2867–2873. 10.1093/bioinformatics/btq559

Miga, K. H., Koren, S., Rhie, A., Vollger, M. R., Gershman, A., Bzikadze, A., Brooks, S., Howe, E., Porubsky, D., Logsdon, G. A., Schneider, V. A., Potapova, T., Wood, J., Chow, W., Armstrong, J., Fredrickson, J., Pak, E., Tigyi, K., Kremitzki, M., … Phillippy, A. M. (2020). Telomere-to-telomere assembly of a complete human X chromosome. Nature, 585(7823), 79–84. 10.1038/s41586-020-2547-7

Pinto, B. J., O'Connor, B., Schatz, M. C., Zarate, S., & Wilson, M. A. (2023). Concerning the eXclusion in human genomics: The choice of sex chromosome representation in the human genome drastically affects the number of identified variants. G3: Genes|Genomes|Genetics, 13(10), jkad169. 10.1093/g3journal/jkad169

Raudsepp, T., Das, P. J., Avila, F., & Chowdhary, B. P. (2012). The pseudoautosomal region and sex chromosome aneuploidies in domestic species. Sexual Development: Genetics, Molecular Biology, Evolution, Endocrinology, Embryology, and Pathology of Sex Determination and Differentiation, 6(1–3), 72–83. 10.1159/000330627

Wang, Z., Sun, L., & Paterson, A. D. (2022). Major sex differences in allele frequencies for X chromosomal variants in both the 1000 Genomes Project and gnomAD. PLoS Genetics, 18(5), e1010231. 10.1371/journal.pgen.1010231

Webster, T. H., Couse, M., Grande, B. M., Karlins, E., Phung, T. N., Richmond, P. A., Whitford, W., & Wilson, M. A. (2019). Identifying, understanding, and correcting technical artifacts on the sex chromosomes in next-generation sequencing data. GigaScience, 8(7), giz074. 10.1093/gigascience/giz074

Zook, J. M., Catoe, D., McDaniel, J., Vang, L., Spies, N., Sidow, A., Weng, Z., Liu, Y., Mason, C. E., Alexander, N., Henaff, E., McIntyre, A. B. R., Chandramohan, D., Chen, F., Jaeger, E., Moshrefi, A., Pham, K., Stedman, W., Liang, T., … Salit, M. (2016). Extensive sequencing of seven human genomes to characterize benchmark reference materials. Scientific Data, 3(1), 160025. 10.1038/sdata.2016.25

